# Volitional and forced running ability in mice lacking intact primary motor cortex

**DOI:** 10.1101/2025.05.14.653913

**Authors:** Ryusei Abo, Mei Ishikawa, Rio Shinohara, Takayuki Michikawa, Itaru Imayoshi

## Abstract

The coordination of various brain regions achieves both volitional and forced motor control, but the role of the primary motor cortex in proficient running motor control remains unclear. This study trained mice to run at high performance (>10,000 rotations per day or >2,700 rotations per hour) using a running wheel, and then assessed the effects of the removal of bilateral cortical areas including the primary motor cortex on volitional and forced running locomotion. The control sham-operated group revealed a quick recovery of volitional running, reaching half of the maximum daily rotation in 3.9 +/-2.6 days (n = 10). In contrast, the cortical injury group took significantly a longer period (7.0 +/-3.3 days, n = 15) to reach half of the maximum volitional daily rotation, but recovered to preoperative levels in about two weeks. Furthermore, even 3 days after surgery to remove cortical regions, the running time on a treadmill moving at 35.3 cm/sec, which is difficult for naïve mice to run on, was not significantly different from that in the sham-operated group. These results suggest that the intact primary motor cortex is not necessarily required to execute trained fast-running locomotion, but rather contributes to the spontaneity of running in mice.

## Introduction

Even seemingly simple behaviors such as reaching out, grasping an object or walking are achieved through a high degree of coordination between various brain regions. Although the rhythms of walking are thought to be produced by a neural circuit in the spinal cord called the central pattern generator (CPG) (Brown, 1911; Grillner and El Manira, 2020), the role of the cerebral motor cortex in the gait control has been the subject of various studies. Canines and felines with cooled motor cortex could walk on flat surfaces but have difficulty walking on ladders and wire mesh, suggesting that the motor cortex is involved in the coordination and adaptation of skilled walking movements rather than in basic gait rhythm generation (Armstrong, 1986). Feline motor cortex neurons change their activity in accordance with the phase of gait (Beloozerova and Sirota, 1993), and electrical stimulation of motor cortex neurons has been shown to alter hindlimb movements during walking (Bretzner and Drew, 2005). In mice, spontaneous walking and grooming movements have been reported immediately after damage to the bilateral cortical primary motor cortex (Nicholas and Yttri, 2024). Unilateral ischemic strokes in the motor cortex of the mouse resulted in slight and transitory deficits in the skilled placement of the contralateral fore paw and hind paw without affecting fundamental aspects of locomotion during swimming, walking and wading (Zorner et al., 2010). After traumatic injury to the unilateral mouse sensorimotor cortex, forced walking or running movement on a treadmill appeared normal, and detailed kinetic analysis has only reported minute effects (Ueno and Yamashita, 2011). These reports have in common that the brain mechanisms of gait motor control depend to a small extent on the cortical motor cortex. Still, the difference in the role of the motor cortex in executing volitional and forced running locomotion remains elusive.

Spinning wheel locomotion has been observed in the wild and reported to be part of play, escape and exploratory behavior (Meijer and Robbers, 2014). Spontaneous wheel running is less stressful and closer to mice’s natural locomotor pattern than forced treadmill exercise (Manzanares et al., 2018). Furthermore, running wheels have been used as a model for exercise training, as the number and frequency of wheel rotations can be electronically recorded and analyzed for individual data (Goh and Ladiges, 2015). On the other hand, treadmill testing does not rely on voluntary activity and can be completed in a short time, making it suitable for elucidating the mechanisms involved in motor adaptation and qualitatively assessing walking movements (Dougherty et al., 2016; Schmitt et al., 2020).

In this study, mice were trained using a running wheel to acquire the locomotor functions that enable high-speed running. Then, the effects of cortical damage on both volitional and forced high-speed running ability were assessed using a running wheel and treadmill, respectively (Fig. 1A). The results elucidated a novel aspect of the role of the primary motor cortex in trained gait motor control.

**Figure 1.**
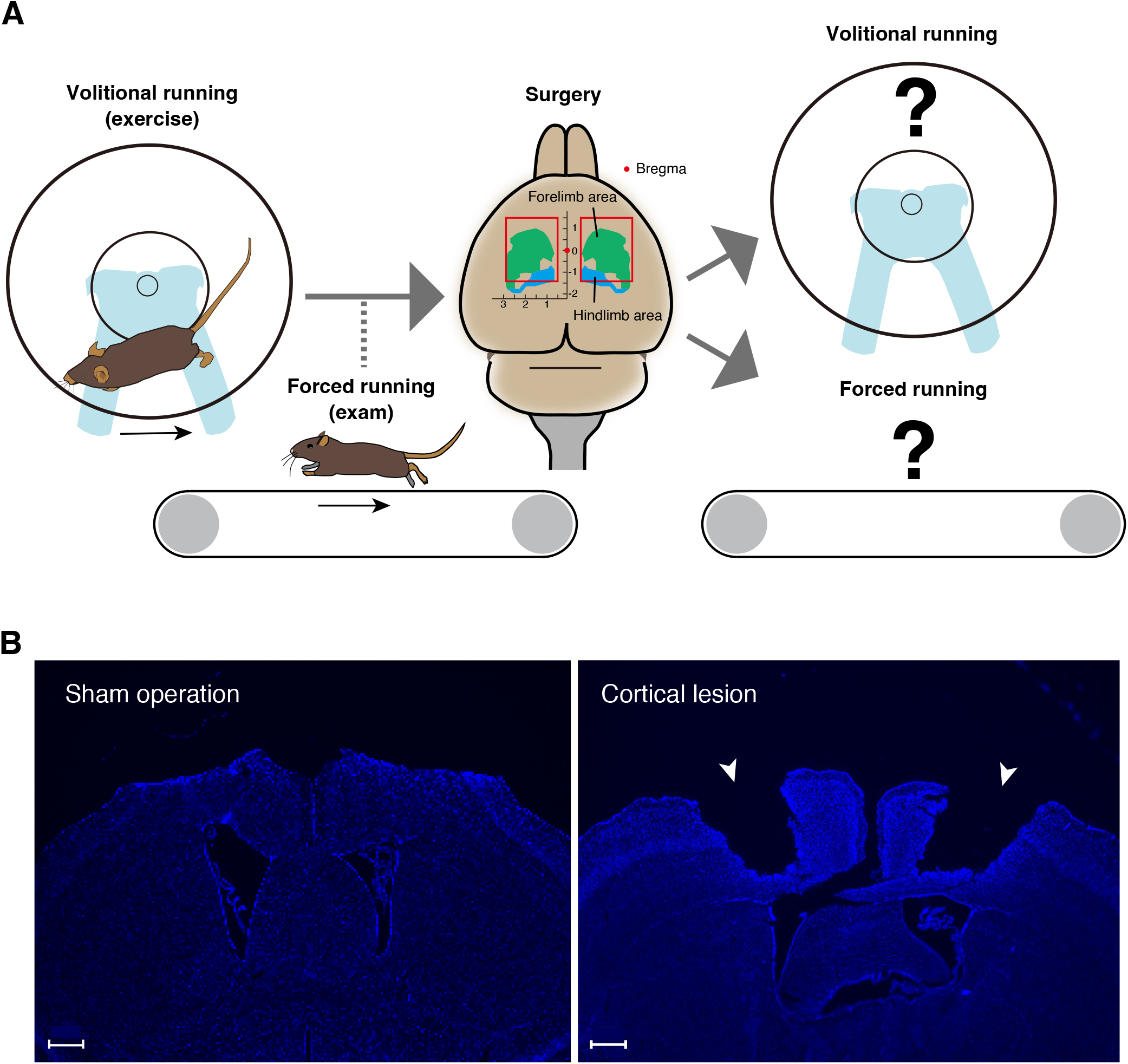
Conceptual diagram of the research strategy. (A) A running wheel is introduced in the home cage to encourage mice to perform volitional running movements; once the total number of rotations per day has reached the criteria, running ability is measured on a treadmill. For mice that have acquired sufficient running ability, surgery is performed to aspirate the bilateral cortical areas including primary motor cortex. Volitional running with a running wheel and forced running on a treadmill are then measured. (B) DAPI staining images of the cerebral cortical section (0.38 mm from the bregma) of the mouse two weeks after sham operation in the left panel and the section (−0.34 mm from the bregma) of the mouse two weeks after injury in the right panel. Injured sites are indicated by arrowheads. Scale bars: 500 µm.

## Materials and Methods

### Animals

Forty-four wild-type C57BL/6J mice (CLEA Japan Inc. and Japan SLC Inc.) at 12-15 weeks of age were subjected to cortical injury surgery, in which the neocortical primary motor cortex was removed by aspiration. Seventeen mice were subjected to sham surgery, in which the skull was detached and then put back together without cortical injury. Fifteen of the 22 mice that survived after cortical injury (8 males and 7 females) and 11 of the 14 mice that survived after sham surgery (4 male and 7 females) were used in the running wheel experiments. Five cortical injury mice (4 males and 1 female) and 4 sham-operated mice (1 male and 3 females) were used in the treadmill experiments.

Mice were delivered to the breeding racks in the behavioral laboratory building one week prior to the experiment. Individuals were identified by ear holes. Mice were monitored for health and body weight and were housed in Innocage (W234×D373×H140 mm) (Innovive) in a temperature- and humidity-controlled room with a 12-hour light-dark cycle reversed (dark period from 8 am to 8 pm, light period from 8 pm to 8 am the following morning). The cages were covered with floor matting and fed pelleted food and water in bottles ad libitum. The treadmill forced running experiment was conducted during the dark period, and intervention on the mice by non-test subjects was avoided as much as possible during the experiment.

### Surgery

After 12-to 15-week-old mice were anesthetized with a triad of anesthetics (medetomidine hydrochloride: 0.0225 mg/mouse, midazolam: 0.12 mg/mouse, butorphanol tartrate 0.15 mg/mouse), the scalp was cut along the midline and the primary motor cortex (anteroposterior axis direction with bregma as origin: - 1.25 ∼ +1.50 mm, left-right axis: 0.50 ∼ 2.75 mm, depth: 0 ∼ 1.00 mm) over the cranium, which was removed with an electric drill to remove bone and the dura mater was stripped to expose tissue. Exposed cortical tissue was removed using an aspirator fitted with a 1 mm diameter 200 µl tip until the tissue was light in color and close to the corpus callosum. The aspirated primary motor cortical area was filled with Spongel (LTL Pharma) moistened with saline to stop bleeding and the removed bone was returned. This procedure was performed bilaterally. The open scalp of the mice was then sutured and the three anesthetic antagonists atipamezole (0.0225 mg/mouse), the antibiotic carprofen (0.2 mg/mouse) and dexamethasone (0.4 mg/mouse) were administered intraperitoneally.

A control group was also prepared in which only tissue was exposed without injury and then sutured (sham-operated group). In the sham-operated group, the exposed brain surface was kept moist with saline so that the operation time was similar to that of the cortical injury group, and the skull was returned and the scalp sutured after being left for a time equivalent to the 15 minutes for the suction operation in the cortical aspiration group.

### Measurement of volitional running locomotion using a running wheel

Battery-powered wheel rotation counter were constructed as described previously (Terstege and Epp, 2024). A running wheel (LIFKOME, 18 cm diameter) was introduced into the mouse cage daily from 12-19 days before surgery. To avoid the influence of light on the diurnal rhythm, battery replacement and data acquisition were carried out in the dark during the dark period. The number of rotations was measured for 24 or 22 consecutive hours per day.

### Measurement of forced running ability using a treadmill

The experiment using the treadmill was based on the method of Dougherty et al (Dougherty et al., 2016) and was performed as follows. This experiment was performed during the dark periods of the day before, 3 days, 1 week and 4 weeks after surgery. The treadmill (MISUMI, SVKR-150-400-6NV-NH-NM-H-R) was darkened by placing a ceiling at the end of the running direction, and the rest of the treadmill was covered with transparent acrylic panels to prevent jumping out. The mice were placed on a treadmill that did not move for the first 60 seconds. Over the next 60 seconds, the mice were allowed to run while increasing their speed from 4.4 cm/sec to 17.7 cm/sec, and then on the treadmill moving at 17.7 cm/sec for the next 60 seconds. The running direction was instantly switched and the mice were allowed to run at 17.7 cm/sec for 60 seconds in the same manner. The speed was increased from 17.7 cm/sec to 35.3 cm/sec in 200 seconds and finally 35.3 cm/sec for 300 seconds. If the mice were unable to run midway, the treadmill was stopped and the mice were retrieved. The upper half of the direction of travel of the mice on the treadmill was covered with a top plate. Under conditions requiring a high speed of 35.3 cm/sec and sufficient running ability of the mice, the mice predominantly traveled in a dark area. On the other hand, when the mouse is temporarily stationary, the mouse appears in a bright area without a top plate. If this condition persists and running becomes impossible, the mouse hits the wall behind it. In order to quantify the running function of the mouse, the total time the mouse was hidden by the top plate when the treadmill was driven at 35.3 cm/sec was measured.

### Fluorescence staining

Mice were deeply anesthetized with a triad of anesthetics followed by perfusion fixation in phosphate-buffered saline (PBS) and 4% paraformaldehyde (PFA)/PBS (pH 7.4). Brains were fixed overnight at 4°C in 4% PFA and replaced every 24 hours with 10%, 20% or 30% sucrose/PBS. After replacement was completed, brains were embedded in an embedding agent (OCT compound, Tissue-Tek) for frozen tissue section preparation and frozen at -80°C. Brain sections (50 µm thick) were prepared in a cryostat (Leica CM1950).

For fluorescence staining, frozen sections were incubated with 5% normal donkey serum and 0.1% Triton X-100/PBS for 1 hour at room temperature for blocking. Afterwards, the sections were washed with PBS and stained with DAPI (4’,6-diamidino-2-phenylindole, Invitrogen, #D21490) (dilution ratio: 1:5000). Sections were attached to glass slides and photographed under a fluorescence microscope (KEYENCE, BZ-X800) (objective: x2, NA = 0.1, #BZ-PA02; filter: BZ-X filter DAPI, excitation: 360/40 nm; emission: 460/50 nm, #OP-87762).

### Data analysis

To determine the extent of recovery of volitional running locomotion after surgery, the maximum number of rotations per day in each mouse was determined by fitting a sigmoid function to the total number of rotations per day and the number of days it takes to reach half of that value was calculated. The sigm_fit function (https://jp.mathworks.com/matlabcentral/fileexchange/42641-sigm_fit) was used to fit the sigmoid function. Student’s t-test was used to compare the number of days it takes to reach half of the maximum wheel rotations between the sham-operated and cortical injury groups. Statistical analysis of the maximum number of rotations per day in volitional running wheel running was performed using the multiple comparison method with the Tukey-Kramer method with the multicompare function in the Statistics and Machine Learning Toolbox of MATLAB, followed by a one-factor analysis of variance (ANOVA). For the treadmill data analysis, the time that mice were in the dark was measured using a custom-made MATLAB code and multiple comparisons were performed using the Tukey-Kramer method.

## RESULTS

### Measurements of volitional running before and after cortical injury

Based on the structure of the motor cortex of C57BL/6 mice, which was determined from the measurement of movements evoked by intracortical micro-stimulation and the histological structure of the cortex (Tennant et al., 2011), the motor cortex including the forelimb and hindlimb areas was removed by aspiration (the anteroposterior axis direction: -1.25 ∼ +1.50 mm, the left-right axis direction: 0.50 ∼ 2.75 mm, and the depth: 0 ∼ 1.00 mm with bregma as the origin) (Fig. 1A). As a control group, a ‘sham operation group’ was created in which the same procedure was performed up to skull and dura mater removal, and the skull was returned and scalp sutured without aspiration of the cortex. Mouse brain sections with sham operation and cortical injury surgery stained by DAPI are shown in Fig. 1B.

To investigate the effects of cortical damage, the amount of volitional locomotion of mice in the home cage was measured using a running wheel capable of recording the number of rotations. An electrical circuit using a microcontroller board (Arduino UNO R3), a magnetic sensor KY-003 and a real-time clock module DS3231 (Terstege and Epp, 2024) was used to record wheel rotation time. To record total daily movements in the home cage, the rechargeable AA batteries were changed once a day, and the mice’s runs were recorded continuously day after day.

The total number of wheel rotations per day of naïve mice, i.e. mice with no prior running-wheel experience, is shown in Fig. 2A. Naïve mice aged 12∼13 weeks took 3.8 days to reach half of the maximum number of rotations per day and about a week to reach the maximum (Fig. 2A). Mice with a plateau in running performance (> 10,000 rotations per day and/or > 2,700 rotations per hour) underwent cortical injury surgery or sham operation. The sham-operated group, in which the scalp was sutured without aspirating the cortex after craniotomy, showed a quick recovery, reaching half of the maximum daily rotation in 3.9 +/-2.6 days (n = 10) (Fig. 2D). In contrast, the cortical injury group took 7.0 +/-3.3 days (n = 15) to reach half of the maximum daily rotation, which is significantly longer than those of the sham-operated group (Fig. 2D). However, both groups recovered their maximal daily wheel rotations to preoperative levels (naïve: 22,801 +/-6,065 rotations/day [n=16]; sham: 22,613 +/-3,511 rotations/day [n = 6]; lesion: 22,885 +/-4,719 rotations/day [n=10]; Fig. 2E).

**Figure 2.**
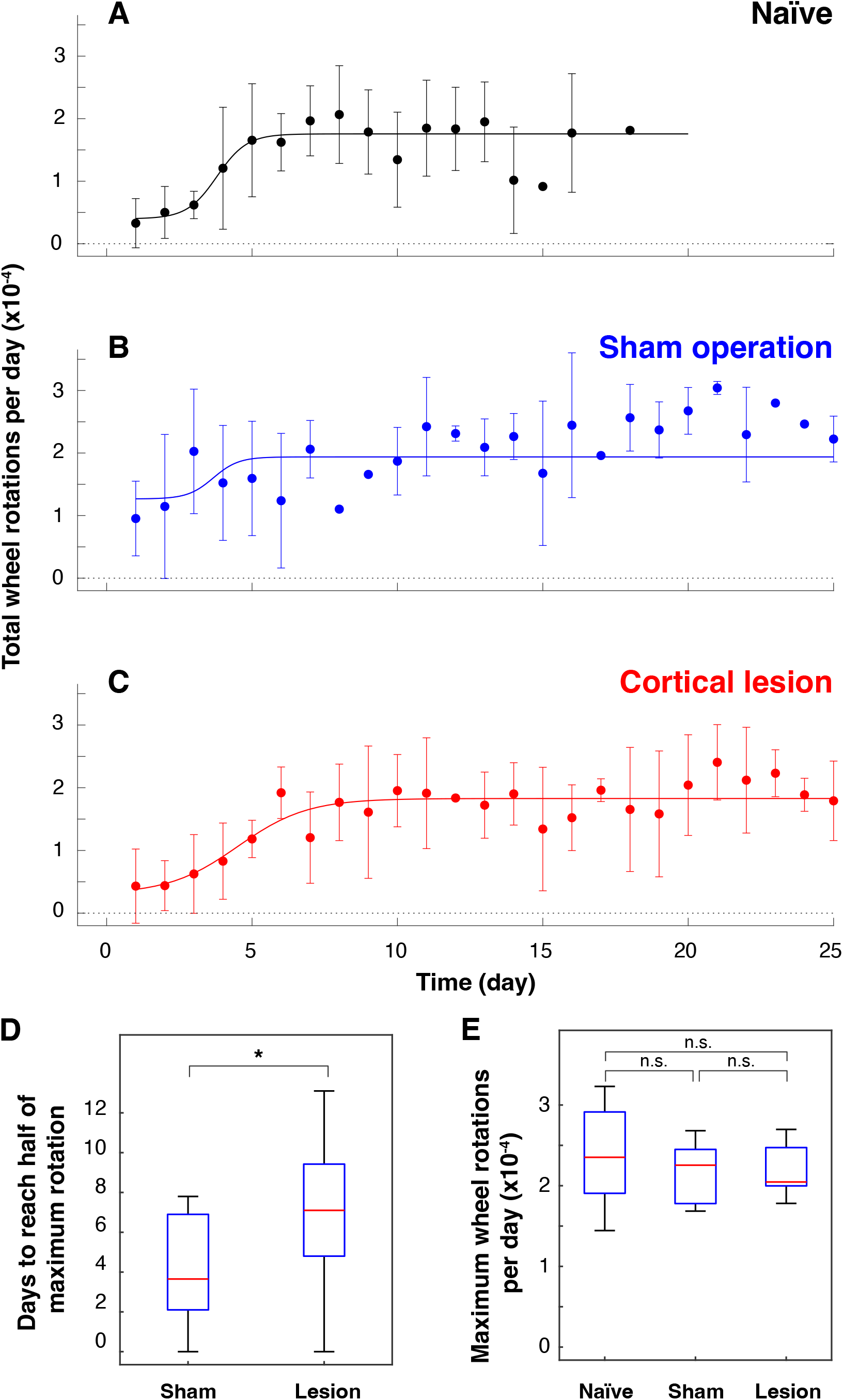
Effects of cortical damage on volitional locomotion in mice. Total wheel rotations per day before and after surgery plotted against elapsed days (mean +/-standard deviation). (A) before surgery (n = 16), (B) sham surgery group (n = 6), (C) cortical injury group (n = 10); (D) number of days taken to reach half of maximum wheel rotations of sham surgery group (n = 10) and cortical injury group (n = 15). * p = 0.0226 (Student’s t-test); (E) maximum number of rotations per day (n. s.: not significant, Tukey-Kramer’s multiple comparison). The results of 24-hour measurements are shown in A-C and E, and the results of both 22- and 24-hour measurements are shown in D (see Materials and Methods).

### Measurements of forced running ability before and after cortical injury

The above volitional running performance measurements showed that the period of reduced running performance after surgery was longer in the cortical injury group than in the sham-operated group. However, locomotion measurements using a running wheel may be influenced by factors other than the ability to perform locomotion, such as motivation to exercise. Therefore, the effect of cortical damage on running motor function was investigated in an experiment in which the subjects were forced to perform running exercise on a rotating treadmill. The details of the experimental procedures for the treadmill running task are described in the Experimental Methods section. Nine of the 14 naïve mice could not run at all on the 35.3 cm/sec treadmill (Fig. 3). Mice trained on a running wheel and achieving running ability beyond the certain standard (> 10,000 rotations per day and/or > 2,700 rotations per hour) were all able to run for more than 180 seconds out of 300 seconds on a treadmill moving at 35.3 cm/sec (Fig. 3). In this study, these high-performing mice were subjected to cortical injury surgery or sham operation. The running times of these mice when the treadmill was driven at a speed of 35.3 cm/sec were compared at preoperative, 3 days, 1 week and 4 weeks after surgery (Fig. 3). Comparison of running times showed no significant differences between all groups between the cortical injury group and the sham operation group (multiple comparison method using the Tukey-Kramer method) (Fig. 3). These results indicate that the ability to run at high speeds was not compromised by cortical damage.

**Figure 3.**
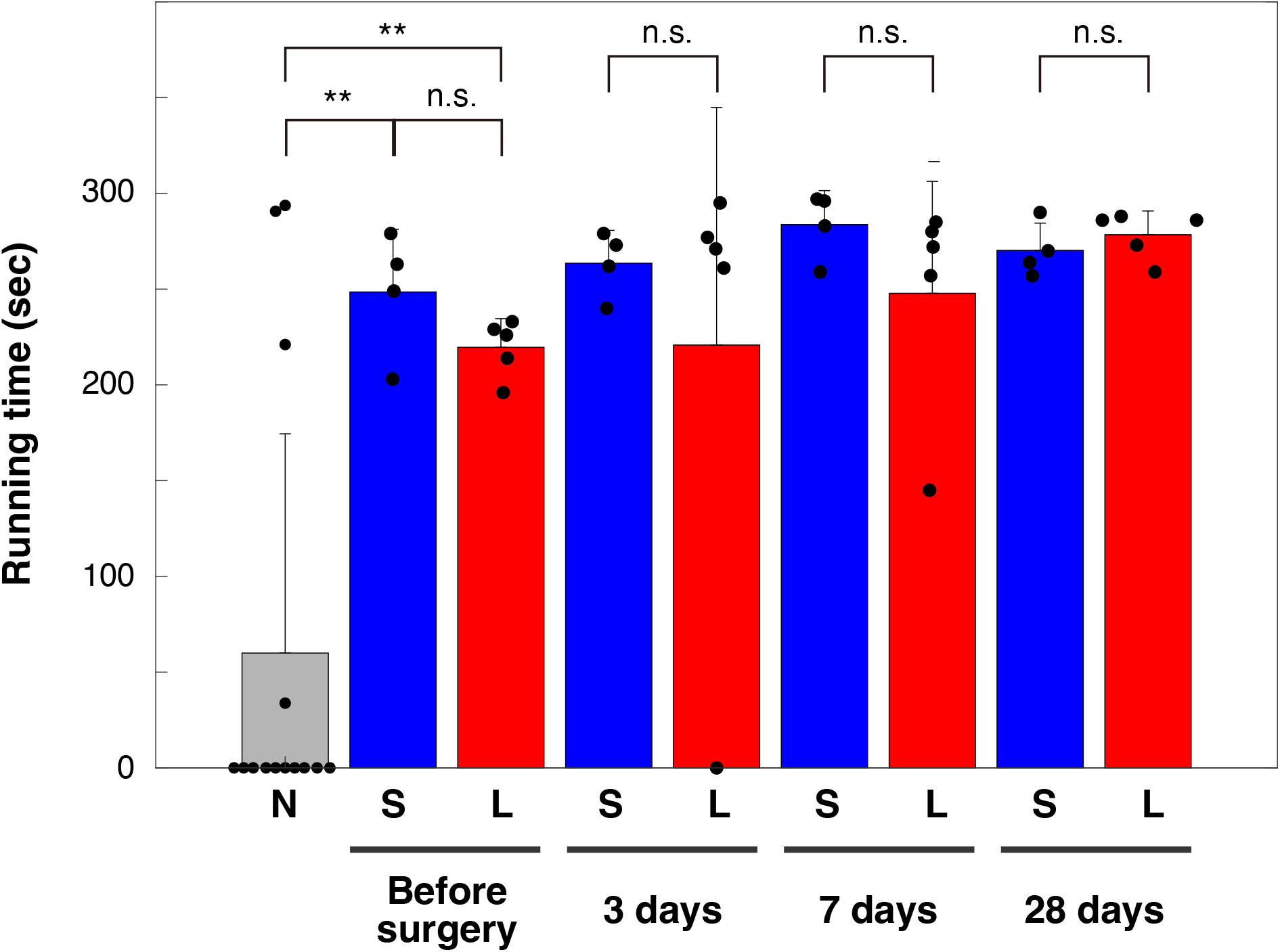
Running time on a treadmill moving at 35.3 cm/s. Bars are means, error bars are standard deviations. Individual values are indicated by black circles. N: naïve mice without running wheel exercise (n = 14), S: shame-operation group (n = 4), L: cortical lesion group (n = 5). **: p< 0.01, n. s.: not significant, Tukey-Kramer’s multiple comparison.

## Discussion

The maximum number of rotations per hour for the mice used was approximately 3,000 when wheel rotation was monitored using the method described previously (Terstege and Epp, 2024). This ability to run sustained over an hour or more can only be reached by training for a week (Fig. 2A). In the present study, mice that were once able to perform running locomotion at this level were operated on by removing the bilateral cerebral cortex including the primary motor cortex, and the effect on running locomotion ability was investigated. It was found that in the sham-operated group, the total number of wheel rotations recovered to the pre-surgery level quickly after surgery. In contrast, in the cortical injury group, it took about 10 days after surgery. On the other hand, when the ability to run was measured on a treadmill moving at a speed of 35.3 cm/sec, which is difficult for naïve mice to run on, there was no significant difference in the time they could run compared to mice in the sham-operated group, even 3 days after cortical injury. These results suggest that the intact primary motor cortex is not necessarily required for the execution of trained running locomotion in mice, which is consistent with previous studies (Armstrong, 1986; Zorner et al., 2010; Ueno and Yamashita, 2011; Nicholas and Yttri, 2024). It has been shown that motor learning involves synaptic plasticity in pyramidal cells of the primary motor cortex (Fu et al., 2012; Xu et al., 2009; Yang et al., 2009). There may be a mechanism by which memories related to motor learning are transferred to other brain regions outside the primary motor cortex during gait motor mastery.

The mesencephalic locomotor region (MLR) (Orlovsky et al., 1999) is located in and near the mesencephalic and pedipalp bridge capsular nuclei where continuous micro-electrical stimulation induces walking movements on a treadmill. The MLR, which projects to the nucleus reticularis gigantocellularis and nucleus reticularis magnocellularis and contributes to the activation of the spinal gait CPG. The basal ganglia constantly inhibits the MLR and the disinhibition initiates walking movements (Roseberry et al., 2016). As the basal ganglia receive excitatory input from the cerebral cortex (Arber and Costa, 2022; Gomez-Ocadiz and Silberberg, 2023; Shepherd, 2013), the cortex controls basal ganglia activity by thought to be involved in the initiation and cessation of gait locomotion. However, so far it is not clear which areas of cortical activity are involved in the initiation and cessation of walking movements. The results of this study suggest that the primary motor cortex may contribute to volitional gait initiation. This is because it took significantly longer for the cortical injury group to reach half of their daily maximum volitional rotation than the sham-operated group, even though the former’s forced running ability was indistinguishable from that of the latter. However, the injured group eventually recovered to preoperative levels of maximum daily wheel rotations, suggesting that there may be neural circuits that can initiate movement even when the intact primary motor cortex is missing. Further investigations of the relationship between activity in areas other than the primary motor cortex, such as the posterior parietal cortex (Drew and Marigold, 2015), and walking movements will help to elucidate the brain mechanisms that control the timing of gait initiation and cessation.

## Ethics statement

All experiments in this study were carried out after approval (Lif-K24014) by the Kyoto University Animal Experimentation Committee and in compliance with animal experimentation guidelines.

## Conflict of Interest

The authors declare that the research was conducted in the absence of any commercial or financial relationships that could be construed as a potential conflict of interest.

## Author Contributions

RA: Investigation, Writing – original draft. MI: Investigation; RS: Investigation; II: Supervision, Writing - review & editing; TM: Conceptualization, Supervision, Investigation, Writing – original draft, Writing – review and editing.

## Funding

This work was supported by JSPS KAKENHI Grant Number 22K06491 (to T.M.) and Grant-in-Aid for Scientific Research (B) JSPS 21H02485 (to I.I.) from the Ministry of Education, Culture, Sports, Science and the Technology of Japan (MEXT); by LiMe Office of Director’s Research Grants 2023 and 2024 (to T.M.); by Japan Science and Technology Agency (JST), CREST Grant Number JPMJCR2433 (to T.M.), JPMJCR1921 (to I.I.) and JPMJCR1752 (to I.I.); and by the Program for Technological Innovation of Regenerative Medicine Grant 21bm0704060h0001 and 24bm1123049h0001 (to I.I.) and Brain/MINDS Grant 21dm0207090h0003 (to I.I.) and Moonshot Research & Development Program JP22zf0127007 (to I.I.) from the Japanese Agency for Medical research and Development (AMED).

## Acknowledgments

We thank Drs. Hirokazu Tanaka and Keisuke Isobe for their insightful comments on the manuscript. We thank Satoshi Hamada and Drs. Adam T. Guy, Masayuki Sakamoto, Yusuke Suzuki, Mayumi Yamada for their advice.

## Notes

### Competing Interest Statement

The authors have declared no competing interest.

